# Role of the ETV5/p38 signaling axis in aggressive thyroid cancer cells

**DOI:** 10.1101/2025.02.17.637322

**Authors:** Jerry H Houl, Rozita Bagheri-Yarmand, Muthusamy Kunnimalaiyaan, Paola Miranda Mendez, Joseph L Kidd, Ali Dadbin, Andrea Jurado-Ruiz, Parag A Parekh, Ying C Henderson, Nikhil S Chari, Aatish Thennavan, Reid T Powell, Clifford C Stephan, Xiao Zhao, Anastasios Maniakas, Roza Nurieva, Naifa L Busaidy, Maria E Cabanillas, Ramona Dadu, Mark Zafereo, Jennifer R Wang, Stephen Y Lai, Marie-Claude Hofmann

**Affiliations:** Department of Endocrine Neoplasia and Hormonal Disorders, The University of Texas MD Anderson Cancer Center, Houston, TX; Department of Head and Neck Surgery, The University of Texas MD Anderson Cancer Center, Houston, TX; High Throughput Research and Screening Center, Texas A&M Institute of Biosciences and Technology, Houston, TX; Department of Immunology, The University of Texas MD Anderson Cancer Center, Houston, TX; Department of Molecular and Cellular Biology, The University of Texas MD Anderson Cancer Center, Houston, TX; Department of Radiation Oncology, The University of Texas MD Anderson Cancer Center, Houston, TX

**Keywords:** Anaplastic thyroid cancer, MAPK pathway, p38/MAPK14 pathway, ETV5, TP53

## Abstract

Patients with poorly differentiated thyroid cancer (PDTC) and anaplastic thyroid cancer (ATC) face a much poorer prognosis than those with differentiated thyroid cancers. Around 25% of PDTCs and 35% of ATCs carry the *BRAF^V600E^* mutation, which constitutively activates the MAPK pathway, a key driver of cell growth. Although combining BRAF and MEK inhibitors can shrink tumors, resistance often develops. The exact cause of this resistance remains unclear.

We previously found that in PDTC and ATC cells the *BRAF^V600E^*mutation is strongly linked to the expression of ETV5, a transcription factor downstream of the MAPK pathway. In the current study, we observed a significant association between ETV5 expression and the activation of p38, a central component of the MAPK14 pathway. Upon reduction of ETV5 levels, p38 expression and activation decreased, along with its upstream regulators MKK3/MKK6. This suggests that the MAPK and p38/MAPK14 pathways are interconnected and that p38 has oncogenic properties in these cancers.

Using high-throughput screening, we established that combining p38 inhibitors with the BRAF inhibitor dabrafenib showed strong synergy *in vitro*, including in cells resistant to dabrafenib and trametinib that had acquired a secondary *TP53* mutation. We then tested this combination in a genetically engineered mouse model (GEMM) of ATC.

Overall, our findings suggest an oncogenic link between the MAPK and p38/MAPK14 pathways and that combining p38 pathway inhibitors with dabrafenib-targeted therapy could improve treatment outcomes for aggressive thyroid cancers. However, more specific and effective p38 inhibitors are required to fully harness this potential.

## Introduction

Transformation of thyroid follicular cells is associated with early driver mutations in the *BRAF* or *RAS* genes in papillary thyroid cancers (PTCs) and follicular thyroid cancers (FTCs) (1). The accumulation of additional driver mutations, including mutations in the *TERT* promoter, leads to undifferentiated tumors such as poorly differentiated thyroid carcinoma (PDTC) and anaplastic thyroid carcinoma (ATC) (2–5). These rare but highly aggressive malignancies are typically associated with a short median overall survival following diagnosis, unless they are treated with targeted therapy (6,7).

According to our recent study and work from others (3,8), approximately 25% of PDTCs and 35% of ATCs harbor the *BRAF^V600E^* mutation, which causes unrestricted cell proliferation due to constitutive activation of the MAPK signaling cascade (9). The *BRAF^V600E^* mutation also synergizes with *TERT* promoter mutations to increase tumor cell proliferation (10,11). Recently, the survival of patients with BRAF-mutated ATC has been significantly prolonged using neoadjuvant targeted therapies that involve a *BRAF^V600E^* inhibitor (dabrafenib) in combination with a MEK1/2 inhibitor (trametinib) followed by surgical resection (7,12,13). Further, immunotherapy in combination with kinase inhibitors may provide additional survival benefit (14,15). Despite these advances, tumor progression rates remain high, and many patients die early from PDTC and ATC. The molecular mechanisms driving resistance to treatments are poorly understood; however several resistance pathways have been proposed. These include activation of alternate signaling pathways mediated by PI3K/AKT (16), alternate *BRAF* splicing (17,18), or reactivation of the MAPK pathway by receptors of the ERBB or FGF receptor families (19,20), amongst others. Also, genomic heterogeneity of PDTC/ATC cells under drug treatment may stimulate clonal evolution and the development of more aggressive phenotypes (21,22). We recently reported that long-term (over 5 months) exposure of PDTC KTC1 cells (*BRAF^V600E^*) to moderate doses of vemurafenib led to the emergence of additional mutations that not only altered cell morphology but also increased their proliferation rate and triggered protein expression and phosphorylation patterns commonly associated with epithelial-to-mesenchymal transition (16). Interestingly, a distinct subpopulation of these cells developed a *KRAS* mutation (*KRAS^G12D^*) alongside *BRAF^V600E^*, giving them a significant advantage in proliferation and invasion—traits reminiscent of ATC. These findings are clinically important, as we later observed acquired *KRAS^G12D^*, *NRAS^Q61K^*, or *KRAS^G12V^* mutations in 50% of *BRAF^V600E^*PTC and ATC patients during progression on dabrafenib or dabrafenib plus trametinib treatments (23). This highlights a key mechanism in therapy resistance and progression. We and others have also detected acquired secondary mutations in the *TP53*, *PTEN*, *NF1, PIK3CA*, and *NF2* genes (24,25), which are tumor suppressor genes that are so far not efficiently targetable.

Effective therapies for PDTC or ATC patients remain an urgent unmet clinical need that should be addressed due to the high risk of disease progression and poor survival rates. Treatment options are particularly limited in cases with primary or secondary *RAS* and tumor suppressor mutations. We previously identified the E26 transformation-specific (ETS) transcription factor ETV5 as a downstream target of the RAS/MAPK pathway and confirmed its importance for *BRAF^V600E^*-mutated PDTC cell proliferation (26). We also demonstrated its possible role in metastasis due to its ability to drive the expression of *TWIST1*, a known activator of epithelial-to-mesenchymal transition (26).

In the present study, using in silico analysis and pre-clinical models, we identified the stress- and oncokinase p38 (or MAPK14) as a possible downstream target of ETV5. Inhibiting the p38 signaling pathway with pharmacological agents reduced PDTC and ATC tumor cell growth. One of these compounds, ralimetinib, showed synergy with dabrafenib, particularly in cells that were resistant to dabrafenib/trametinib and harbored a secondary *TP53* mutation. This implies that the mutational status of *TP53* affects the response to p38 inhibitors at progression.

## Material and Methods

### Cell lines and culture conditions

We used the human thyroid cancer cell lines MDA-T68, MDA-T85, KTC1, KTC1-VA7, Hth83, PDX.008.CL, and SW1736. We also used the mouse ATC cell lines MCH2.2 and PPA6 (Supplementary Table S1). MDA-T68 and MDA-T85 cells were generated as previously described (27). The MDA-T68 line was derived from a follicular variant of papillary thyroid carcinoma (FVPTC) with cervical lymph node metastasis. This cell line harbors a *NRAS^Q61K^*mutation. The MDA-T85 cell line was derived from a metastatic lymph node in a patient who had a primary PTC with extrathyroidal extension and lympho-vascular invasion. MDA-T85 cells harbor a BRAF*^V600E^* and a *HRAS^Q61K^* mutations as well as the *TERT* promoter mutation C228T, and they are considered PDTC cells. KTC1 are PDTC cells derived from a metastatic pleural effusion (28) and were kindly provided by Dr. R. Schweppe, University of Colorado, Anschutz Medical Center. These cells harbor the *BRAF^V600E^* and the *TERT* promoter C250T mutations (29). KTC1-VA7 is a KTC1 subclone that spontaneously acquired an additional *KRAS^G12D^* mutation after long-term culture with the *BRAF^V600E^* inhibitor vemurafenib (16). Hth83 is an ATC cell line established from a giant cell anaplastic thyroid carcinoma (30). Hth83 cells harbor *HRAS^Q61R^* as a key genetic driver as well as two frameshifts in the *TP53* gene*: TP53^P153Afs*28^* and *TP53^P152fs^* (29). They also exhibit the *TERT C228T* promoter mutation. SW1736 cells are ATC cells harboring a *BRAF^V600E^* mutation, a *TP53^Q192*^* truncating mutation, and the *TERT* promoter *C228T* mutation (29). Hth83 and SW1736 cells were kindly provided by Dr. Jeffrey Myers at the University of Texas MD Anderson Cancer Center. The PDX.008.CL cell line was derived from a mediastinal node metastasis of a dabrafenib-resistant human ATC tumor (31). Mouse cell lines were established from endogenous thyroid tumors developed by genetically engineered mice (*Tpo-Cre-ER^T2^; BRAF^Ca/+^; Trp53^-/-^*, see below). Cell lines were authenticated through short tandem repeat profiling (STR) either by the MD Anderson Cytogenetic Core facility (human cell lines) or by IDEXX BioAnalytics (North Grafton, MA, USA) (mouse cell lines). All lines were routinely tested for *Mycoplasma* by PCR. Cells were maintained in RPMI growth media supplemented with glutamine, pyruvate, penicillin/streptomycin, and 10% fetal bovine serum. All reagents were purchased from Thermo Scientific (Waltham, MA, USA).

### Long-term cultures for testing resistance

Separate dishes of MDA-T85 cells were cultured long-term for 32 passages (32 weeks/8 months) in the continuous presence of 0.03% DMSO (control) or 5 μM dabrafenib in 0.03% DMSO in RPMI media. Cell subpopulations were labeled MTA-T85.R0 (control) and MTA-T85.R2 to MTA-T85.R8 according to the number of months kept in long-term cultures. The overall passage number status of these subpopulations of cells following the long-term dabrafenib treatment was p40 from their original line derivation. Following the long-term dabrafenib treatment, the genetic identity of the resistant cell subclones was confirmed using STR fingerprinting. IC_50_ for dabrafenib in these subclones was measured by culturing the cells at 0.0, 0.01, 0.1, 1.0, 5 and 10 μM of the drug in 0.1% v/v DMSO. Three biological replicates were performed for each experiment. A nontreated plate was fixed and nuclei were stained with 4′,6-diamidino-2-phenylindole (DAPI) at the time of drug treatment (Day 0) to provide the number of cells present at the start of treatment as previously described (32). After 72 hours of incubation in the presence of drug or 0.03% v/v DMSO (negative control), plates were fixed with 0.4% paraformaldehyde and nuclei stained with DAPI using an integrated HydroSpeed plate washer (Tecan Life Sciences) and Multidrop Combi dispenser. Plates were imaged on an IN Cell Analyzer 6000 laser-based confocal imaging platform (General Electric Healthcare Bio-Sciences) and nuclei counted using the algorithms developed using the IN Cell Developer Toolbox software (version 1.6). Dabrafenib and trametinib resistance were confirmed by using real-time live cell imaging of T85-R0 and T85-R4 cell growth in an IncuCyte® S3 System (Sartorius, Bohemia, NY, USA). The cells were plated in 96-well plates in triplicates and imaged for 72 hours in the presence of either dabrafenib or trametinib at the indicated concentrations.

### DNA sequencing

Whole exome sequencing (WES) of DNA samples from MDA-T85 cells treated long-term with 5 µM dabrafenib as described above and collected at month 0 (R0), month 2 (R2), month 4 (R4) and month 8 (R8) was performed by the MD Anderson Cancer Center Advanced Technology Genomics Core (ATGC). For comparison, DNA samples were collected from MDA-T85 cells treated long-term with 0.03% DMSO and collected at month 0, month 2, month 4, and month 8 of treatment.

The BWA-MEM algorithm was used to align the raw sequencing reads with the human reference genome (GRCh38). The resulting alignments were processed to generate sorted BAM files. Duplicate reads were marked using Picard MarkDuplicates. Base quality score recalibration was performed using GATK BaseRecalibrator. Somatic single nucleotide variants (SNVs) and small insertions/deletions (indels) were called using GATK Mutect2. Dabrafenib-resistant and matched DMSO BAM files were jointly analyzed to identify new somatic mutations. Variant filtration was applied using GATK FilterMutectCalls to remove low-quality calls. The variants were annotated using Funcotator.

Copy number alterations were detected from the WES data using GATK ModelSegments. Segmented copy number data were then analyzed using GISTIC2.0 to identify significantly amplified and deleted regions across samples. GISTIC was run using a q-value threshold of 0.25 and a confidence level of 0.95 to identify driver copy number events.

### Mouse model and mouse cell lines

Mouse studies were approved by the University of Texas MD Anderson Cancer Center Institutional Animal Care and Use Committee (IACUC) and conducted in compliance with the NIH Guide for the Care and Use of Laboratory Animals and the IACUC-approved protocols. All mice were obtained from the Jackson Laboratory. An ATC transgenic mouse model was generated as described by McFadden et al (33). Briefly, a *Tpo-Cre-ER^T2^; BRAF^Ca/+^; Trp53^fl/fl^* transgenic strain was created. Upon administration of tamoxifen in 2 months old mice, the animals developed thyroid tumors after a latency time of 3-4 months, as expected. Tumor growth was followed using MRI, then tumors excised after animal euthanasia. For histopathology, tumor fragments were fixed in Bouin’s fixative, dehydrated using increasing ethanol concentrations and xylene, and then embedded in paraffin. After deparaffinization and rehydration, 5 µm tissue sections were prepared and stained with hematoxylin and eosin. Histological analysis demonstrated an evident pattern of undifferentiated cells resembling human ATC, with small clusters of PTC cells in one of seven tumors analyzed (Supplementary Figures 1A and 1B). Five cell lines were derived from 18 different ATC mice and characterized (for genotypes of some lines, see Supplementary Figure 1C). Only two cell lines, MCH2.2 and PPA-6, were used in this study. Genotypes before (mouse tails) and after *Cre* recombination (tumors and cell lines) were assessed with the appropriate primer pairs (https://jacks-lab.mit.edu/protocols). Mouse ATC cell lines were cultured in RPMI growth media supplemented with glutamine, pyruvate, penicillin/streptomycin, and 10% fetal bovine serum (all reagents from Thermo Fisher Scientific).

### Analyses of publicly available datasets

Gene expression values of ETV5, p38, and MEF2A comparing human ATC cells to normal thyroid cells were obtained from publicly available databases (NCBI-GEO GSE65144 (34) and GSE33630 (35)). Data were extracted from 23 ATC and 57 normal thyroid tissue samples. The probe expression matrices we downloaded had been preprocessed and standardized. Correlations between ETV5 expression, p38 expression, and expression of components of the MAPK14 pathway were obtained from The Cancer Genome Atlas (TCGA) using cBioPortal (1).

### Gene downregulation with shRNA and validation/quantification

Lentivirus particles containing ETV5-shRNA or control shRNA (scrambled) were used to infect MDA-T85 cells (MISSION^®^ shRNA, Sigma-Aldrich, St Louis, MO, USA). Transfected cells were selected in media containing 2 µg/mL puromycin (Clontech/Takara Bio, Mountain View, CA, USA). The expression of ETV5 and two known ETV5 targets (TWIST1, SNAI1, Supplementary Table S2) were quantified by quantitative real-time PCR (qRT-PCR) to validate the downregulation of ETV5 expression.

### Quantitative real-time PCR

Total RNA was isolated using the Zymed Laboratories kit (Thermo Fisher Scientific, Waltham, MA, USA). cDNA was produced using the SuperScript VILO master MIX (Thermo Fisher Scientific, Waltham, MA, USA). Quantitative real-time PCR (qRT-PCR) was performed using TaqMan probes from Applied Biosystems (Thermo Fisher Scientific, Waltham, MA, USA) (Supplementary Table S2). All analyses were performed using previously described methods (31). In brief, the expression value of each gene was normalized to the expression of an internal control gene (GAPDH or ACTB, ΔCt), and then compared to the experimental controls using the ΔΔCt method (fold change). qRT-PCR was performed using three technical replicates per sample and experiment, along with a minimum of three independent experiments. The *p*-values ≤ 0.05 were considered significant.

### Western blot analysis and quantification

Cells were lysed in a buffer composed of 50□mM HEPES, pH 7.5, 1% (vol/vol) Igepal-C630, 1□mM EDTA, 10□mM NaF, 1□mM sodium orthovanadate, 5% (vol/vol) glycerol, 50□mM NaCl, and 1□mM PMSF (36). Halt Protease Inhibitor Cocktail and Phosphatase Inhibitor Cocktail were added at standard concentrations. All reagents were purchased from Thermo Fisher Scientific, Waltham, MA, USA. Lysates were boiled at 100°C for 5 minutes, briefly centrifuged, and the supernatant loaded onto 10 % polyacrylamide or 4% to 12% Stain-Free TGX gels (5-25 µg/lane) (Bio-Rad, Hercules, CA, USA). After separation, proteins were transferred onto nitrocellulose or PVDF membranes with a Trans-Blot Turbo Transfer System (Bio-Rad, Hercules, CA, USA). Membranes were blocked with 5% BSA in Tris buffer saline with 0.1% Tween 20 (TBST) for 60 minutes, or EveryBlot Blocking Buffer (Bio-Rad, Hercules, CA, USA) for 5 min. The blots were incubated with antibodies recognizing human ETV5, human p38, human phospho-p38, or additional components of the p38 signaling pathway (Supplementary Table S3). Anti-ACTB, anti-Vinculin, or anti-GAPDH were used as loading controls (Supplementary Table S3). Blots were incubated with primary antibodies at 4°C overnight. After washing with TBST, the membranes were incubated with the appropriate horseradish peroxidase-conjugated secondary antibodies (Supplementary Table S3) for 1 hour at room temperature. Blots were washed and incubated with chemiluminescence substrates (SuperSignal West Dura #34095 and SuperSignal™ West Femto #34095, Thermo Fisher Scientific, Waltham, MA, USA) and visualized with a ChemiDoc MP Imager imaging system (Bio-Rad, Hercules, CA, USA). Protein band intensities were quantified using ImageJ software (NIH) or Image Lab 6.1 (Bio-Rad).

### High-throughput drug screening

High-throughput screens were performed by the Gulf Coast Consortia’s (GCC) Combinatorial Drug Discovery Program at Texas A&M Health Science Center Institute of Bioscience and Technology. Cell lines were screened against dabrafenib (a *BRAF^V600E^* inhibitor), several p38 kinase inhibitors (ralimetinib, PD169316, and SB203580), a putative TWIST1 inhibitor (harmine), a BCL2 inhibitor (venetoclax), a PARP1/2 inhibitor (olaparib), a prenylated chalconoid and NOTCH inhibitor (xanthohumol), and the cytotoxic agents anysomycin and doxorubicin (positive controls). The cell lines MDA-T68 (PTC, with a dominant follicular component) and Hth83 (ATC) were used as negative controls as they do not harbor any BRAF mutation. For screening assays, a total of 500 cells in 50 µL of RPMI media were seeded in each well of black high binding uClear® 384-well polystyrene plates (Greiner Bio-One, Monroe, NC, USA) using a Multidrop Combi liquid dispenser (Thermo Fisher Scientific, Waltham, MA, USA). After seeding, the plates were incubated at room temperature for 45-60 min then placed into a humidified cell culture incubator with 5% CO_2_ at 37°C. The cells were allowed to form a monolayer overnight, then 50 nL of drugs at different concentrations were transferred into each well using an Echo 550 acoustic dispensing platform (Labcyte/Beckman Coulter, Indianapolis, IN, USA). The drugs were tested at a final concentration of 0.00, 0.01, 0.10, and 1.00 μM in 0.1% v/v DMSO. Two biological replicates were performed for each experiment. A nontreated plate was fixed and nuclei were stained with 4′,6-diamidino-2-phenylindole (DAPI) to determine the number of cells present at the start of drug treatment (Day 0) as previously described (32). After 72 hours of incubation in the presence of drug or 0.1% v/v DMSO (negative control), plates were fixed with 0.4% paraformaldehyde and nuclei stained with DAPI using an integrated HydroSpeed plate washer (Tecan Life Sciences, Morgan Hill, CA, USA) and Multidrop Combi dispenser. Plates were imaged on an IN Cell Analyzer 6000 laser-based confocal imaging platform (General Electric Healthcare Bio-Sciences), and nuclei were counted using the algorithms developed using the IN Cell Developer Toolbox software (version 1.6) (32).

Statistical evaluation was performed on each plate throughout the course of the screening to assess the consistency of results as previously described (32). Metrics evaluated included the rate of growth of the negative controls, the coefficient of variance of the positive and negative controls, and assay robustness determined from the *z*-scores. Assay reproducibility and experimental drift were determined as previously reported (32). We used the normalized growth rate inhibition algorithm which quantifies the cellular response to drugs (37).

### High-throughput screening combination of anchor and probes

We performed a high-throughput combination screen using an anchor-probe strategy on six cell lines (MDA-T85, MDA-T68, Hth83, SW1736, MCH2.2 and PPA-6). Dabrafenib, a BRAF^V600E^ inhibitor, was selected as the anchor drug because of its FDA-approved use in ATC patients (7). Dabrafenib was combined with seven inhibitor compounds that target pathways identified from public databases or array analyses (26). Those inhibitors were p38 inhibitors (ralimetinib, PD169316, SB203580), harmine (a putative TWIST1 inhibitor), venetoclax (a BCL2 inhibitor), and xanthohumol (a prenylated chalconoid). Eight concentrations of probe molecules were tested in pairwise combinations with five concentrations of the anchor drug (100, 55, 30, 20, and 10 nmol/L). Drug interactions were categorized as synergistic, additive or antagonistic by comparing the experimentally observed dose-response surface to a theoretical surface calculated from the independent single-agent activity using a Bliss synergy model as previously described (32). As a cutoff, we used the sum of all pairwise interactions to define antagonism (< −1) or synergy (> +1), respectively. Additional drug synergy metrics, including excess over the highest single agent, Loewe additivity, and zero interaction potency, were calculated using the SynergyFinder Plus R package. In addition to providing expanded metrics to quantify drug synergy, this package also uses a bootstrapping method to evaluate the statistical significance of the drug-drug interaction, thus providing a robust evaluation of drug-drug synergy.

### Animal treatments

For a preliminary assessment, syngeneic mice (equal numbers of males and females) were subcutaneously (flank) injected with approximately 1 × 10^6^ MCH2.2 mouse ATC cells. Tumor growth was measured using a caliper. Treatments were initiated once the tumors achieved a volume of 100 mm^3^, about 10 days after cell injection. The mice were randomly divided into treatment groups of five mice each: vehicle control (0.01% DMSO in phosphate buffered saline), dabrafenib (BRAF^V600E^ inhibitor), ralimetinib (p38/MAPK14 inhibitor), PD169313 (p38/MAPK14 inhibitor), SB203580 (p38/MAPK14 inhibitor), harmine (TWIST1 inhibitor), and xanthohumol (NOTCH inhibitor). All reagents were prepared at a final concentration of 30 mg/kg in 0.01% DMSO and injected as single treatments. Tumor size was measured and recorded for another 12 days, and all mice were euthanized at that point. A Kaplan-Meier survival graph for the different treatments was established.

For drug combination studies with mice bearing endogenous tumors, *TPO-Cre-ER^T2^; BRAF^Ca/+^; Trp53^fl/fl^* transgenic mice were treated with tamoxifen using standard procedures (Jackson Laboratory). After three months, tumor volumes were regularly measured using MRI until the end of treatments. Once thyroid tumors reached a volume of ∼100 mm^3^, mice (six mice per treatment) were treated with dabrafenib (0, 6, 12, or 30 mg/kg), ralimetinib (0 or 30 mg/kg), or a combination of dabrafenib and ralimetinib (0/0, 6/30, 9/30, 12/30, or 30/30 mg/kg/day respectively). A combination of dabrafenib (6 mg/kg/day) and trametinib (0.6 mg/kg/day) was used as the benchmark. Tumors were measured every 5 days for 21 days using MRI and recorded as percent change in tumor growth compared to tumor volume on day 0 of treatment.

### Statistical analysis

Student *t-*tests or one-way ANOVA were used to compare two or more groups. For data nonparametric data distribution, the Mann–Whitney–Wilcoxon test was used to compare two samples or groups. A *P* value less than 0.05 was considered statistically significant. All statistical tests were two-sided.

### Data Availability

The data generated in this study are available within the article and its supplementary data files.

## Results

### Expression of the transcription factor ETV5 in ATC patient samples and cell lines

We previously demonstrated that the transcription factor ETV5 is expressed downstream of BRAF or RAS mutations in PTC patient samples and the PDTC cell line KTC1 (26). In these cells, ETV5 enhanced growth and acted as a driver of TWIST1 gene expression, which is a well-known mediator of epithelial-to-mesenchymal transition (38). Our GEO databases analyses of ATC patient samples indicated that expression of *ETV5* mRNA and its target *TWIST1* were also significantly upregulated in ATC patient samples compared to normal thyroid tissues (Figure 1A-D). Further, as shown by western blotting, the ETV5 protein was prominently expressed in several PTC and ATC cell lines (Figures 1E and 1F), particularly in the nucleus, as shown by its co-expression with the nuclear marker lamin A (nuclear extracts). ETV5 expression was rapidly down-regulated by BRAF^V600E^ inhibitors in short-term treatments (48 hours) and this effect was dose-dependent. Dabrafenib was more efficient than vemurafenib in the mouse ATC cell line MCH2.2 and a human PDTC cell line (MDA-T85) (Figure 1G). Vemurafenib induced paradoxical ERK1/2 activation, particularly in the MDA-T85 cells, confirming previous findings (39). Furthermore, ETV5 expression was significantly upregulated in a subclone of KTC1 PDTC cells (KTC1-VA7) that acquired a secondary *KRAS^G12D^* mutation upon long-term treatment (5 months) with vemurafenib and became resistant to BRAF^V600E^ inhibitors (38) (Figure 1H).

**Figure 1:**
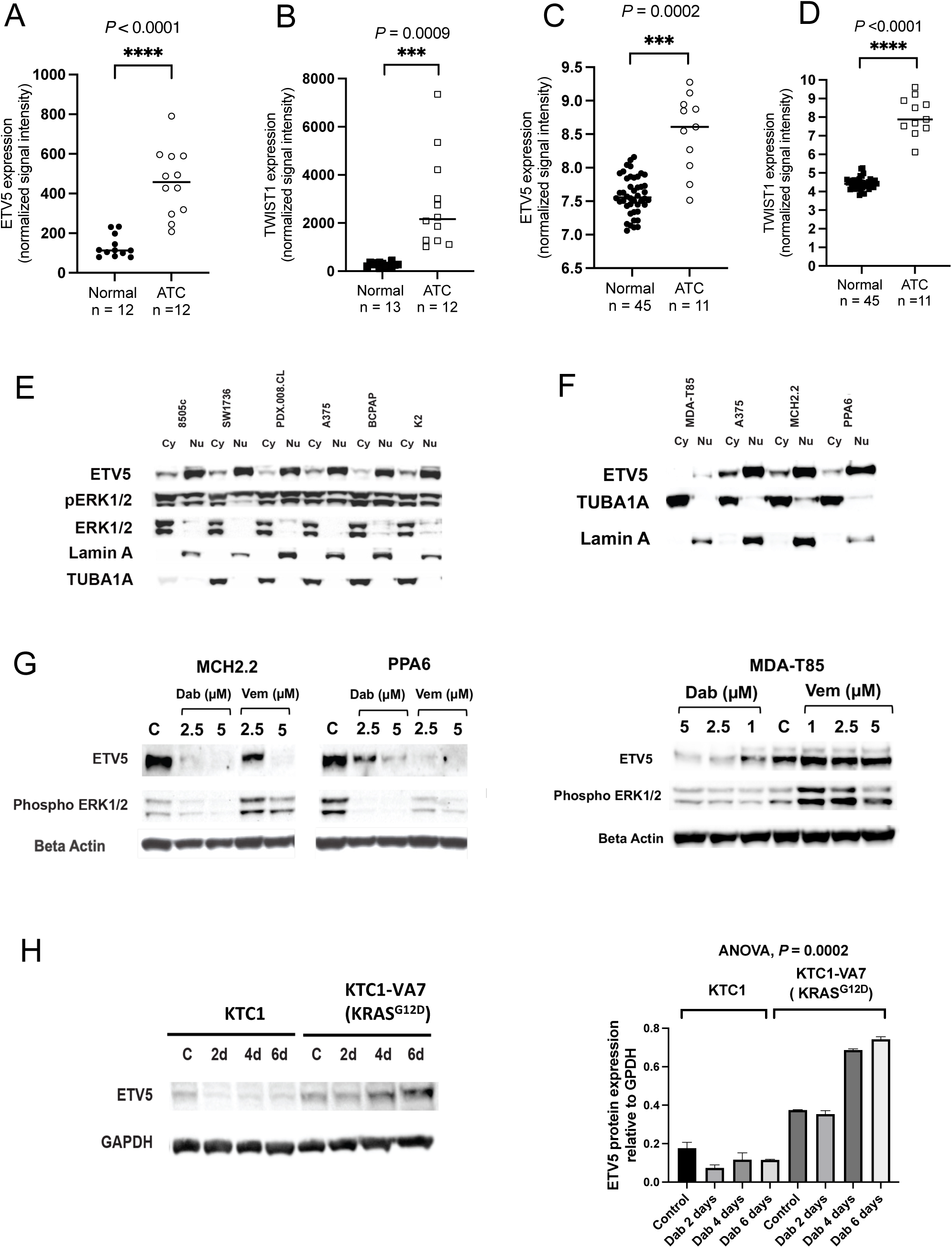
mRNA and protein expression of the transcription factor ETV5 in human ATC samples and cell lines. **A-D**: Analysis of GEO databases for mRNA ETV5 expression and its target TWIST1. **A** and **B**: GSE65144, **C** + **D**: GSE33630. **E:** Protein expression of ETV5 in PTC (BCPAP, K2), ATC (8505c, SW1736, PDX.008.CL), and melanoma (A375) cells. TUBA1A is used as a marker for the cytosol and Lamin A for the nucleus. **F:** Expression of ETV5 in PDTC (MDA-T85), melanoma (A375) and two mouse ATC cell lines (MCH2.2 and PPA6). **G:** Protein expression of ETV5 in two mouse ATC cell lines (MCH2.2 and PPA6) and a human PDTC cell line (MDA-T85) treated with different concentrations of dabrafenib (Dab) or vemurafenib (Vem). ETV5 expression is downregulated in a dose-dependent manner. **H:** Protein expression of ETV5 in the PDTC cell line KTC1 and a BRAF inhibitor-resistant clone of KTC1, KTC1-VA7, after acquisition of a *KRAS^G12D^* mutation (representative western blot and corresponding quantitation, n =3, dabrafenib = 2.0 µM).

### Resistance to dabrafenib in PDTC cells is associated with increased ETV5 and p38 activity

To confirm ETV5 upregulation at resistance, we established another BRAF inhibitor resistant model, whereby MDA-T85 PDTC cells were cultured long-term (up to 8 months) with 5 µM dabrafenib. Cells were assessed every month (R0 to R8) for drug resistance (IC_50_) and expression of ETV5 and ERK1/2 activity. Dabrafenib resistance was demonstrated by an increase in IC_50_ that reached > 50 µM at 4 months of culture (Figure 2A). Interestingly, these cells also became resistant to trametinib and continued to grow compared to drug-sensitive cells despite the high drug concentrations (5-20 µM) (Figures 2B and 2C). We also observed a decrease in ETV5 protein expression during the first months of dabrafenib exposure, followed by a steady recovery over time (Figure 2D). As no specific pharmacological inhibitor of ETV5 is available, we performed an in silico analysis to identify molecules that correlate with ETV5 expression in BRAF-mutated thyroid cancers. We first mined TCGA through cBioportal to analyze mRNA of 496 PTC samples (1). Results indicated that *ETV5* mRNA expression correlated strongly with *MAPK14* (also called *p38*) expression, a kinase often regulated by cellular stress, which also has oncogenic properties (40,41) (Figure 3A). We also observed that p38 expression and phosphorylation/activity followed ETV5 expression over time in dabrafenib-resistant cells (Figure 2D). In addition, expression of *MAP3K2*, a component of the p38MAPK cascade that leads to p38 activation, correlated with that of mutated *BRAF* (Figure 3B). Further, as shown in Figure 3C, we found that *p38* mRNA was significantly upregulated in ATC patient samples compared with normal thyroid tissues. The mRNA expression of *MEF2A*, a target of the activated p38 protein, was also increased in ATC samples (Figure 3D). In addition, p38 protein and its activated/phosphorylated form, as well as ATF2/7, a known target of activated p38, were found in all PDTC and ATC cell lines tested (Supplementary Figure 2).

**Figure 2:**
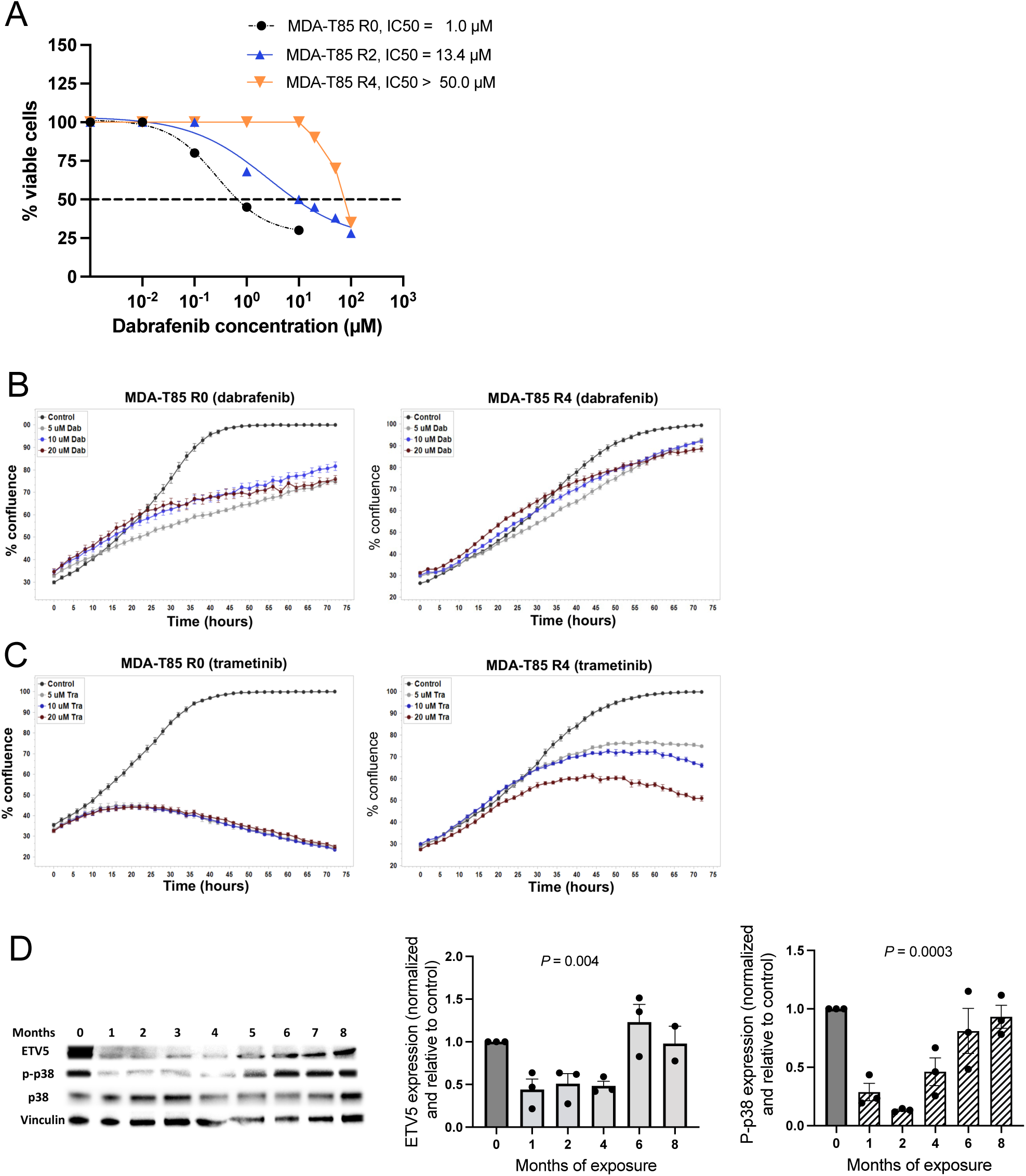
Acquisition of dabrafenib/trametinib resistance and resistance markers in a PDTC cell line (MDA-T85) **A:** Increase of dabrafenib resistance over time measured with the CellTiter-Glo luminescent cell viability assay (R0, R2, R4 = months of dabrafenib exposure, n = 7 for each data point). **B:** Dabrafenib resistance as measured by an increase in cell number (% well confluence after 72h dabrafenib exposure) visualized with the IncuCyte S3 real-time imaging system (n = 3 for each data point ± SEM). **C:** Concomitant trametinib resistance after 72h trametinib exposure similarly quantitated with the IncuCyte S3 system (n = 3 for each data point (± SEM). **D:** Quantitative protein expression of ETV5 and its potential target, phospho-p38, after 1, 2, 4, 6, and 8 months of dabrafenib exposure and acquisition of dabrafenib and trametinib resistance, with representative western blot (n = 3 blots, ± SEM).

**Figure 3:**
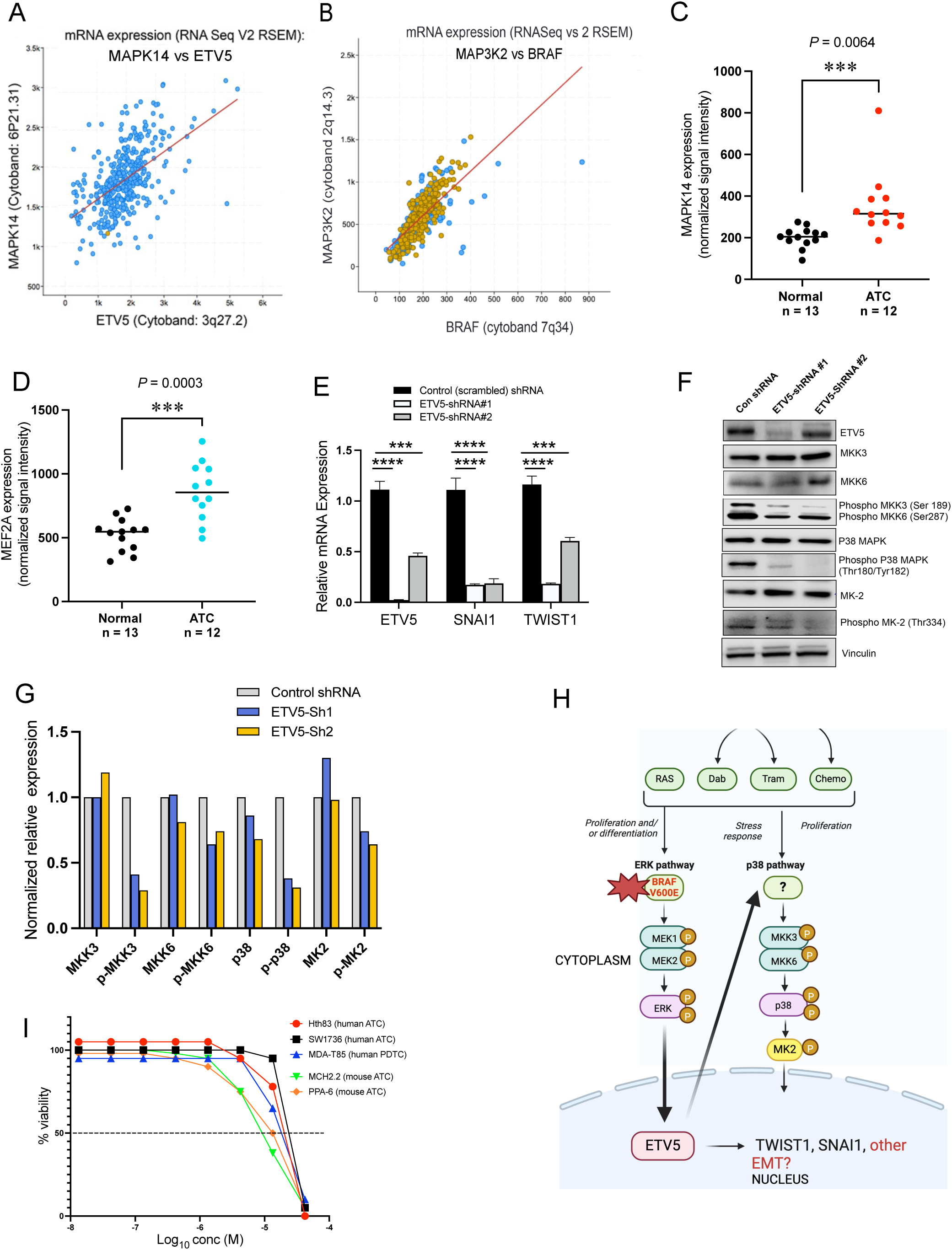
Expression and potential interrelationship between ETV5 and p38 pathway components in PTC/ATC patient samples and PDTC/ATC cells. **A:** ETV5 mRNA expression positively correlates with that of *MAPK14 (p38)* in 397 PTC samples (TCGA/cBioPortal). **B:** Positive correlation between the expression of *BRAF^V600E^* (yellow dots) and *MAP3K2* kinase, an upstream activator of p38 protein activity, in 397 PTC samples (TCGA/cBioPortal). Blue dots = BRAF wild type. **C:** mRNA expression of *MAPK14/p38* in ATC patient samples compared to normal thyroid samples (from NCBI GEO GSE65144). **D:** mRNA expression of *MEF2A*, a substrate for activated p38, in ATC patient samples compared to normal thyroid samples (from NCBI GEO GSE65144). **E:** Quantitative PCR analysis showing down-regulation of *ETV5* mRNA expression in MDA-T85 cell subclones after shRNA treatments. ETV5 downregulation is confirmed through concomitant downregulation of its targets *TWIST1* and *SNAI1*. **F + G:** Down-regulation of ETV5 protein in MDA-T85 cells is associated with downregulation of p38 pathway components. **H:** Potential functional interrelationship between the MAPK and the MAPK14/p38 pathways. **I:** Growth inhibition of different PDTC and ATC cell lines with the p38 pathway inhibitor ralimetinib.

### p38 expression and activity depend on ETV5 expression

To investigate whether ETV5 is responsible for p38 expression/activation, or, conversely, if p38 activity indirectly drives ETV5 expression, we used shRNA to downregulate *ETV5* in MDA-T85 PDTC cells. This down-regulation was confirmed by assessing the expression levels of its targets, *TWIST1* and *SNAI1* (Figure 3E). The observed decrease in both *TWIST1* and *SNAI1* mRNA expression validated the effective inhibition of *ETV5* and provided further support for the involvement of ETV5 in driving epithelial-to-mesenchymal transition (EMT) in these cells (26). Downregulation of *ETV5* expression by shRNA decreased p38 protein expression and inhibited p38 phosphorylation/activity, as observed in Figures 3F and 3G. This observation suggests a functional interrelationship between these two molecules/pathways. Additionally, the reduced phosphorylation of MAPKAPK2 (or MK2), an immediate downstream target of p38 (41), further confirmed the alteration of p38 activity. Downregulation of ETV5 expression also caused a concomitant decrease in the phosphorylation/activity of MKK3/6 kinases. Because these kinases are known triggers of p38 activity (41), our data position ETV5 as a possible upstream regulator of the p38 pathway (Figure 3H). Therefore, through ETV5 upregulation, the MAPK pathway might activate the p38 pathway in thyroid cancer cells. We next assessed the sensitivity of several human and mouse PDTC and ATC cell lines to ralimetinib, a p38 inhibitor (Figure 3I). Our data indicate that the IC_50_ for ralimetinib is 10-13 µM and that the drug equally affects BRAF mutated and BRAF wild-type cell lines.

### High-throughput drug screening identifies synergism between dabrafenib and ralimetinib

Due to the significant occurrence of drug resistance in PDTC and ATC patients and possible cardiovascular adverse events linked to MEK inhibitors (42), it is imperative to explore alternative treatment options that can be combined with dabrafenib. Among these alternatives, we identified various pharmacological inhibitors of p38, including ralimetinib, which has been investigated in phase I trials involving patients with diverse solid tumors and lymphomas, often in conjunction with chemo- and radiotherapies (43–45). Thus, in this study, we investigated the effects of p38 inhibition in combination with dabrafenib in *BRAF^V600E^*-mutated PDTC and ATC cells, and in animal models.

For each cell type, IC_50_ values were determined for each inhibitor (Supplementary Table S4). Dabrafenib IC_50_ values were high (>30 µM) for both BRAF wild-type cell lines (MDA-T68 and Hth83), as expected. The independent single-agent activities were used to calculate theoretical surfaces. Synergy Scores were obtained by comparing the experimentally observed dose response surface to the theoretical surface and were calculated using the difference from Bliss (32,46)(Supplementary Table S5). Drug concentrations of ralimetinib and dabrafenib were categorized as synergistic (green), additive (black) or antagonistic (red) (Figure 4A-4D and Supplementary Table S5). Figures 4A’-4D’ show the corresponding synergy scores for each drug and cell line, with grouping into synergy (>1.00, green), additivity (−1.00 to +1.00, yellow), and antagonism (<-1.00, red). In MDA-T85 cells, which harbor a *BRAF^V600E^*and a *HRAS^Q61K^* mutations, no inhibitor showed significant synergy in combination with dabrafenib (Figure 4A’). Only additivity was detected as well as some antagonistic effects of harmine (TWIST1 inhibitor) and PD169316 (p38 inhibitor). In MDA-T68 cells, which harbor solely *NRAS^Q61K^* as a driver mutation and were used as negative controls, only xanthohumol, a flavonoid with potential NOTCH signaling inhibitory activity, led to significant inhibition. Furthermore, the ATC cell line Hth83, which is BRAF wild type but harbors a *HRAS^Q61R^* and a *TP53^P152fs,P153fs^* mutations, was also used as a negative control and exhibited no synergistic response by any statistical model (Figure 4B’). However, in SW1736, MCH2.2 and PPA-6 cells (Figure 4B-D and 4B’-D’), which all harbor a *BRAF^V600E^* and a *TP53* deletion, p38 inhibitors significantly synergized with dabrafenib. The combination of ralimetinib and dabrafenib exhibited the most consistent response. Since the identity of the ralimetinib target is controversial and could be the EGFR kinase rather than p38 (47), we tested the effect of ralimetinib on the activities/phosphorylation of p38, MK2 (downstream of p38), and EGFR in two cell lines (MTA-T85 and Hth83). In both cell lines, ralimetinib significantly inhibited MK2 phosphorylation, but did not significantly inhibit p38 nor EGFR activity (Supplementary Figure 3). Therefore, in PDTC and ATC cells, ralimetinib inhibited the p38/MAPK14 pathway and decreased cell proliferation (Figure 3I and Supplementary Table S4) but uses a mechanism that differs from its effects in other cancer cell types (47).

**Figure 4:**
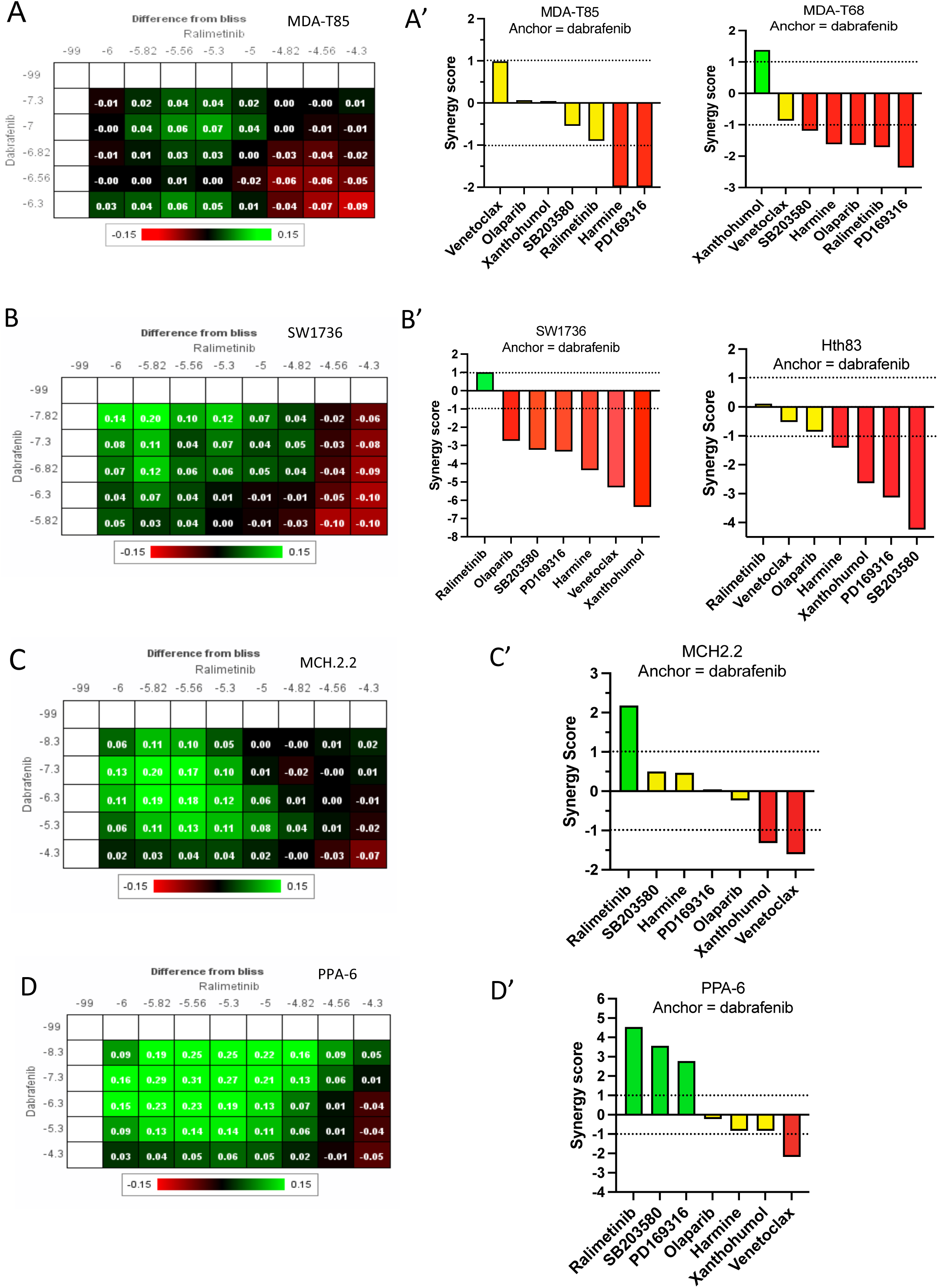
High-throughput analysis indicating synergy between p38 pathway inhibitors and dabrafenib in some PDTC and ATC cell lines. **A-D**: Heat maps showing drug synergy as measured by the growth inhibition difference from that determined using the Bliss independent model. Green areas represent a difference from Bliss >0, indicating synergy. Synergy was observed at multiple drug doses (green areas). Black areas represent additivity and red areas represent antagonism. The drug concentrations (M) are shown on a log scale. **A’-D’**: Synergy scores calculated according to Handley *et al* (37). Green indicates synergy (>1), yellow additivity (≤1) and ≥ −1), and red antagonism (<−1). MDA-T68 (PTC with a dominant follicular component) and Hth83 (ATC) are BRAF wild-type and considered negative controls.

### Ralimetinib synergizes with dabrafenib in dabrafenib-resistant cells

We next assessed how dabrafenib resistance affected the response to pharmacological inhibitors and whether ralimetinib could re-sensitize resistant tumor cells in combination treatments. In MDA-T85 cells, the synergy between dabrafenib and ralimetinib emerged at month 4 (R4) of dabrafenib resistance and persisted over time (Figure 5 and Supplementary Table S6), indicating that inhibition of the p38 pathway can resensitize the cancer cells to dabrafenib, and that p38 might be involved in drug resistance. Our data also indicate that the *TP53* mutation status of the tumor cells (truncation versus gain-of-function, see below) is relevant because *BRAF^V600E^/TP53^-/-^* cells were more sensitive to the combination treatment (dabrafenib 5-15 nM/ralimetinib 1.0-1.5 µM, Figure 4) than *BRAF^V600E^/TP53* gain-of-function cells (dabrafenib 5-10 µM/ralimetinib 15 µM, Figure 5).

**Figure 5:**
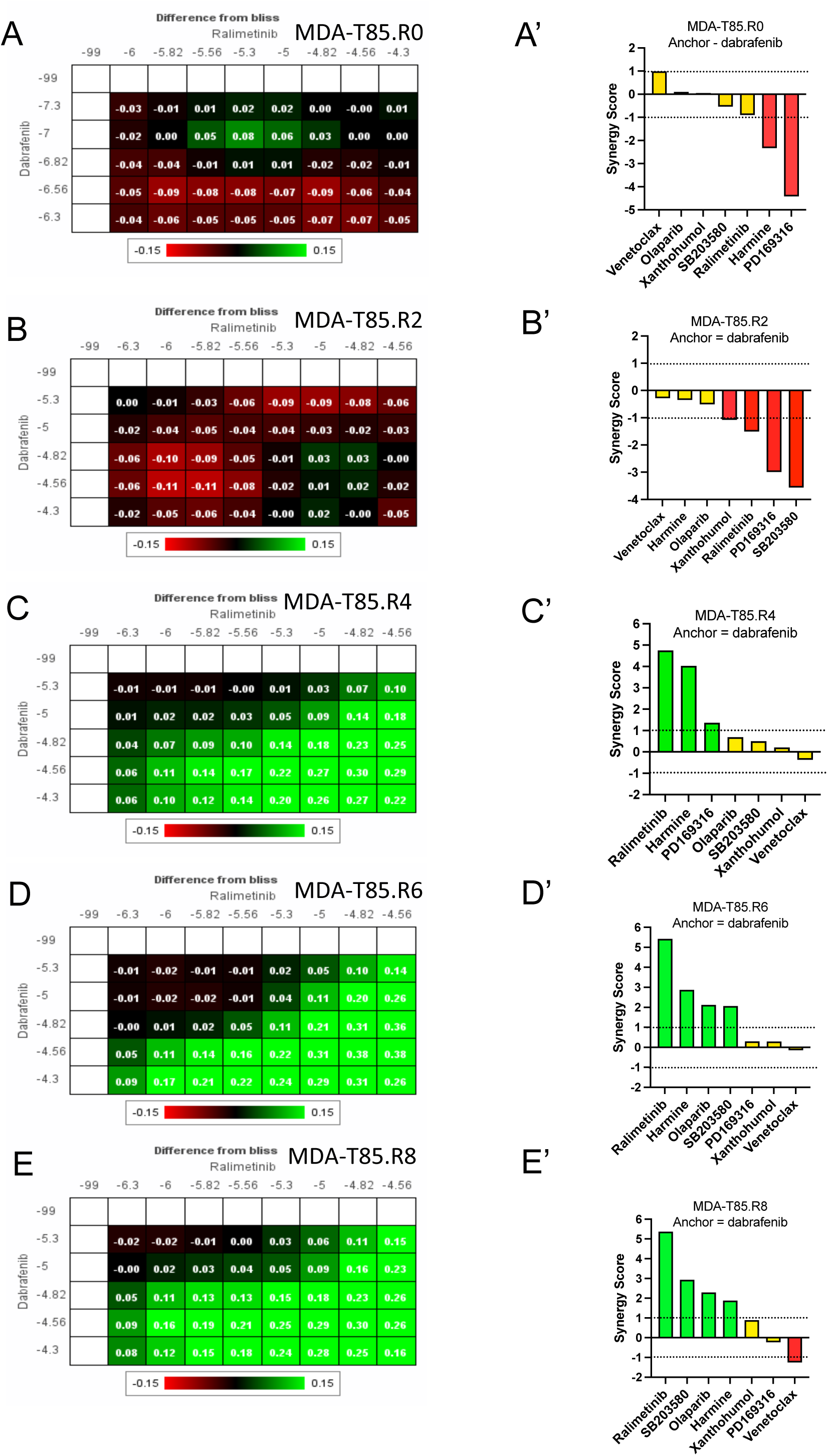
High-throughput analysis indicating synergy between the p38 pathway inhibitor ralimetinib and dabrafenib in dabrafenib/trametinib resistant PDTC cells. *BRAF*^V600E^/*HRAS^Q61K^-* mutated MDA-T85 cells were cultured for 8 months in 5.0 µM dabrafenib or 0.001% DMSO. Cells were harvested each month for further processing, resulting in subclones (R0-R8). R0, R2, R4, R6 and R8 clones were analyzed for drug synergy between dabrafenib and ralimetinib. **A-D**: Heat maps showing drug synergy as measured by the growth inhibition difference from that determined using the Bliss independent model. Green areas represent a difference from Bliss >0, indicating synergy. Synergy was observed at multiple drug doses (green areas). Black areas represent additivity and red areas represent antagonism. The drug concentrations (M) are shown on a log scale. **A’-D’**: Synergy scores calculated according to Handley *et al* (37). Green indicates synergy (>1), yellow additivity (≤1) and ≥ −1), and red antagonism (<−1).

### Resistance to dabrafenib in MDA-T85 cells parallels a *TP53**^R248W^*** driver mutation

Whole exome sequencing of MDA-T85 cells exposed long-term to dabrafenib (sub-clones R0 to R8) revealed a variety of mutations compared to cells exposed to DMSO only. Overall, the number of additional novel mutations increased with the duration of drug exposure (Figure 6A). Most mutations were passenger mutations; however, one documented driver mutation, *TP53^R248W^*, was detected at month 8 of dabrafenib exposure (Figure 6A and Supplementary Table S7). A splice site mutation also occurred in this gene that was deemed not pathogenic. The accumulation of mutations over time indicated an increase of genomic instability possibly driven by dabrafenib. Most mutations were missense mutations (55%), followed by nonsense mutations (7%) and frame shifts/deletions (5%). Most single-nucleotide variants (SNVs) were C>T followed by T>C and C>A and the median SNVs per sample was 15. Finally, GISTIK copy number analysis revealed significant changes mostly at month 8 of dabrafenib exposure (Figure 6B and Supplementary Table S8), with amplifications of chromosome 1p and substantial deletions in chromosome 8 (p+q) and 18 (p+q). Chromosome 1 amplifications are recurring events in thyroid cancers (48) and may be associated with more aggressive or advanced disease states, possibly through amplification of *NRAS* (1p13.2). None of the observed chromosomal deletions affected genes known to drive aggressive thyroid cancers, including at drug resistance (*TP53, PTEN, PIK3CA, ATM, ATR, NF1, NF2, BRCA1/2*).

**Figure 6:**
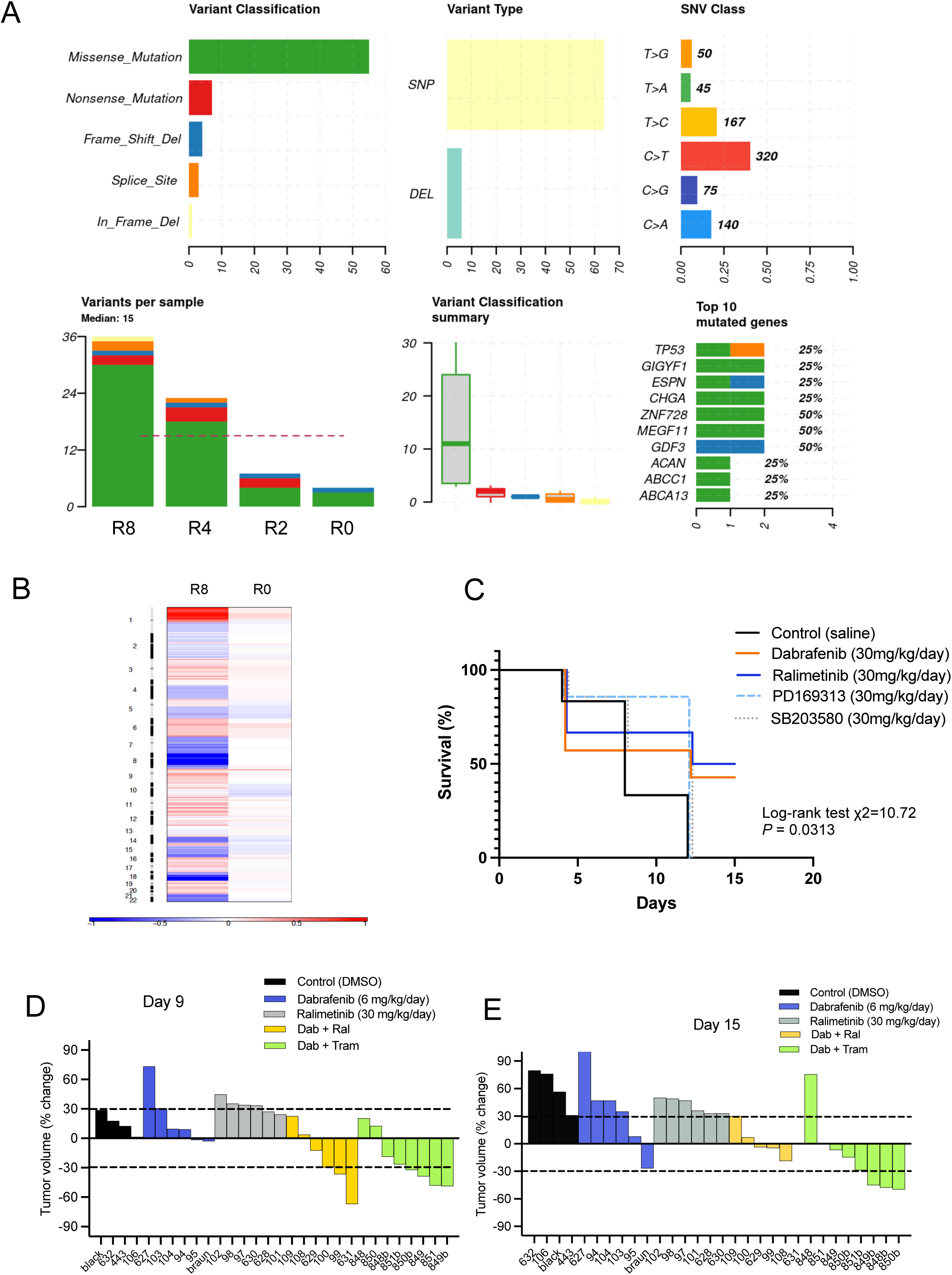
Whole-exome sequencing of dabrafenib/trametinib resistant cell clones and *in vivo* effect of dabrafenib in combination with ralimetinib or trametinib in an ATC mouse model. **A:** GATK somatic mutation analysis indicating accumulation of missense mutations over 8 months of dabrafenib exposure (R0 to R8) and detection of a *TP53* mutation. **B:** GISTIC analysis. Heat map image of MTA-T85 R0 and R8 clones (month 0 and month 8 of dabrafenib exposure). Image was obtained using GISTIC (v2.0.23). Chromosome arrangement flows vertical, top to bottom ordering. Length of chromosomes and their arms are proportionately represented. Red represents copy number gain and blue represents copy number loss in units of log2 (dabrafenib/DMSO treated). Color intensity correlates with the statistical significance of the change. **C:** Kaplan-Meier survival analysis for xenografted mice (endpoint = tumor doubling). Approximately 10^6^ MCH2.2 mouse ATC cells (*BRAF^Ca/+^; p53^del/del^*) were injected subcutaneously into the flank of 30 C57Bl/6 mice. When tumors reached 100 mm^3^, the mice were randomly distributed into the following 5 groups (n=6 per group): Control (saline), dabrafenib, ralimetinib, PD169313, and SB203580. Log-rank test χ2=10.72, *P* = 0.0313. **D+E:** Percentage change in tumor volume for individual ATC mice (waterfall plots). *Braf^+/CA^; p53^flox/flox^* mice were treated with tamoxifen and tumors measured by MRI until volumes reached 100 mm^3^. Tumor-bearing mice were then treated daily with DMSO (0.03% in saline), dabrafenib (6 mg/kg/day), ralimetinib (30 mg/kg/day), dabrafenib (6 mg/kg/day) + ralimetinib (30 mg/ kg/day), or dabrafenib (6 mg/kg/day) + trametinib (0.6 mg/kg/day). D = Measurement at day 9 of treatments, E = measurements at day 15 of treatments. Each color represents a different treatment; each bar represents the % tumor volume change in one mouse. The labels on the x axis represent the mice ID tags.

### Ralimetinib combined with dabrafenib shows some synergy in ATC mouse models

To assess the effect of p38/MAPK inhibitors in ATC mouse models, we first tested these drugs by xenografting MCH2.2 mouse tumor cells in syngeneic C57Bl/6 mice (flank injections). Mice were euthanized when tumors reached a volume of 100% of initial volume. Ralimetinib and dabrafenib yielded similar responses (∼50% of mice survived beyond day 15 of treatment) (Figure 6C) both at the dose of 30 mg/kg/day. Other p38 inhibitors (PD169313 and SB203580) had only transient effects and these mice were euthanized at day 12 of treatment (median survival = 12 days), as were the control animals (median survival = 8 days).

We then assessed different doses of dabrafenib in combination with trametinib (dabrafenib 0, 6, 12 and 30 mg/kg/day; trametinib 0 and 0.6 mg/kg/day) in *Braf^+/CA^; Trp53^del/del^* ATC mice. Tumor volumes were assessed using MRI. Data indicated that, in these mice, dabrafenib (6 or 12 mg/kg/day) in combination with trametinib (0.6 mg/kg/day) significantly reduced tumor growth compared to the control treatment (DMSO only). Therefore, to assess synergy, the lower dabrafenib dose (6 mg/kg/day) was tested alone and in combination with ralimetinib or trametinib. During the first 9 days of treatment, the dabrafenib-ralimetinib combination was as effective as the dabrafenib-trametinib combination but this effect did not persist as smaller tumor volume reductions were detected on day 15 and beyond (Figures 6D and 6E).

## Discussion

In this study, we identified the p38/MAPK14 pathway as a downstream target of ETV5 in PDTC and ATC preclinical models. Pharmacological inhibitors of p38 signaling reduced tumor cell growth and showed synergistic effects with dabrafenib. Our data also indicated that the mutational status of *TP53* might affect the response to p38 inhibitors, particularly in dabrafenib-resistant cells.

The ETS family of transcription factors contains 28 homologs that can be categorized into 12 subfamilies (including ETS and PEA3) based on their sequence similarity and the positioning of the ETS domain (49,50). These transcription factors play crucial roles in various biological processes, including tumor progression (50). As shown in the present study, ETV5 was also expressed in multiple aggressive thyroid cancer cell lines (PDTC and ATC). Its expression was associated with activation of the MAPK pathway and was downregulated by dabrafenib, corroborating our previous data (26). Furthermore, ETV5 was re-expressed at drug resistance, making it a potential marker for tumor progression. Because there is no drug that effectively inhibits ETV5 nor any PEA3 transcription factors, we focused on inhibiting one of its putative downstream targets, p38, which is also re-expressed at resistance.

p38 plays a key role in the cellular response to stress and inflammation. It is essential for maintaining cellular homeostasis and mediating various signaling pathways, including p53. Dysregulation of p38 has been linked to several diseases including inflammatory disorders, neurodegenerative diseases, and cancer progression (41,51). In our study, ralimetinib was the most effective of the p38 inhibitors tested, but there is recent controversy regarding the identity of its target (47). As demonstrated here, ralimetinib effectively downregulated the p38 pathway by specifically inhibiting the kinase activity of MK2, a direct downstream target of p38, but not EGFR kinase activity. The observed synergy between dabrafenib and ralimetinib *in vitro*, and to some extent *in vivo*, supports the need for further preclinical investigations using p38 inhibitors with better specificity and efficiency.

Acquired, or secondary, resistance often develops after an initial period of response to kinase inhibitors, including combination therapies such as dabrafenib and trametinib. The mechanisms of drug resistance vary extensively among individuals and tumor types, and identifying these mechanisms should improve the clinical efficacy of treatments. Known mechanisms of resistance include target alteration, which reduces drug binding or increases kinase activity (52,53); novel gene driver mutations (16,23,31); bypass signaling that often activates the PI3K/AKT pathway (16,54–57); and growth factor receptors’ amplification (19,31,58). Some of these resistance mechanisms have been observed using cell lines derived from melanoma, hepatocarcinoma, and colon cancer in addition to patient samples. In the present study, we used a BRAF inhibitor resistance model of aggressive thyroid cancer cells. Resistance to dabrafenib appeared between months 3 and 4 of drug exposure and was accompanied by resistance to trametinib as well. These features are similar to the clinical observations (40) and our previous data obtained with another PDTC cell line (16). In the absence of additional genomic mutations in mediators of the MAPK or PI3K-AKT pathway, the cause of this concomitant resistance to trametinib might be the epigenetic upregulation of receptors such as EGFR1 (ERBB1) and ERBB3 (19), known for activating both the PI3K-AKT and MAPK pathways (59). This would also explain the rebound of ETV5 and concomitant p38 upregulation at resistance.

In BRAF-mutated solid tumors, common causes of resistance to dabrafenib and trametinib include *KRAS* or *NRAS* mutations (60). It is unknown whether selective pressure favors the development of pre-existing mutated subclones or whether BRAF inhibition itself induces genomic instability over time. In the present study, confirming previous data (16), the secondary driver mutation was not detectable by WES at the start of treatment nor in the parental cells. *TP53^R248W^* is a hotspot gain-of-function mutation that occurs in the DNA-binding domain of the p53 protein. This mutation is one of the most commonly mutations observed in cancers, particularly in those arising in the breast and head and neck regions, where it is more potent in driving tumorigenesis and metastasis compared to other *TP53* mutations (61,62). Similar to other mutant p53 proteins, mutant p53^R248W^ undergoes strong Hsp90-mediated stabilization, leading to the accumulation of high protein levels in the nucleus and cytoplasm (63). Gain-of-function mutant p53 proteins interact with and significantly impact various signaling pathways, including the STAT3 and p38 pathways (64,65). In the present study, ralimetinib and its synergistic activity with dabrafenib were more efficient when p53 protein expression was deleted rather than overexpressed by a gain-of-function *TP53^R248W^*secondary mutation. The indirect enhancement of MKK3/6 phosphorylation by both ETV5 and gain-of-function p53 could potentially shift p38 function toward a more aggressive oncogenic role (64,66,67), but further investigations are needed to understand this crosstalk in the context of PDTC/ATC. Also, a transcriptomic assessment of clonal evolution over time will help elucidate evolutionary patterns and shifts in gene expression associated with dabrafenib resistance.

In summary, ETV5, p38, and p53 play interconnected and critical roles in the regulation of PDTC/ATC cell behavior. ETV5, often associated with enhanced proliferation, migration, and metastasis in cancer, is tightly linked to signaling pathways such as p38/MAPK, which can exhibit dual roles (apoptosis or tumorigenesis) depending on the cellular context. The interaction between p38 and p53 mutation status adds further complexity. Dysregulation of this network can shift the balance between tumor suppression and growth, highlighting their combined importance in cancer biology. We found that, in these cells, ralimetinib inhibited the p38 pathway by targeting MK2 activity, and significantly synergized with dabrafenib *in vitro* in the presence of *TP53* mutations. Ralimetinib also synergized with dabrafenib during the initial days of ATC mice treatments. Despite the lack of sustained *in vivo* effects, our work has highlighted a novel resistance pathway and positioned ETV5 and p38 as key markers of tumor progression. Targeting the intricate crosstalk between these factors using improved p38, HSP90, or STAT3 inhibitors (68–70), possibly combined with immunotherapy, offers a promising avenue for developing more effective therapeutic strategies for these tumors.

## Acknowledgements

This work was supported by the Anaplastic Thyroid Cancer Multidisciplinary Research Funds (JHH, RBY, YCH, AD, SYL, MCH); a University of Texas MD Anderson Cancer Center IRG Seed Grant (MCH); and DOD #HT9425-23-1-0675 (MCH). We also thank the Institute for Biotechnology (IBT) High Throughput Research and Screening Center at Houston (CPRIT grant number RP200668) and acknowledge the NIH/NCI for supporting the MD Anderson Cancer Center Core Facilities (NCI P30CA016672). Editorial support was provided by Bryan F Tutt, Senior Scientific Editor, Research Medical Library at MD Anderson Cancer Center.

---

**Supplementary Figure 1.**
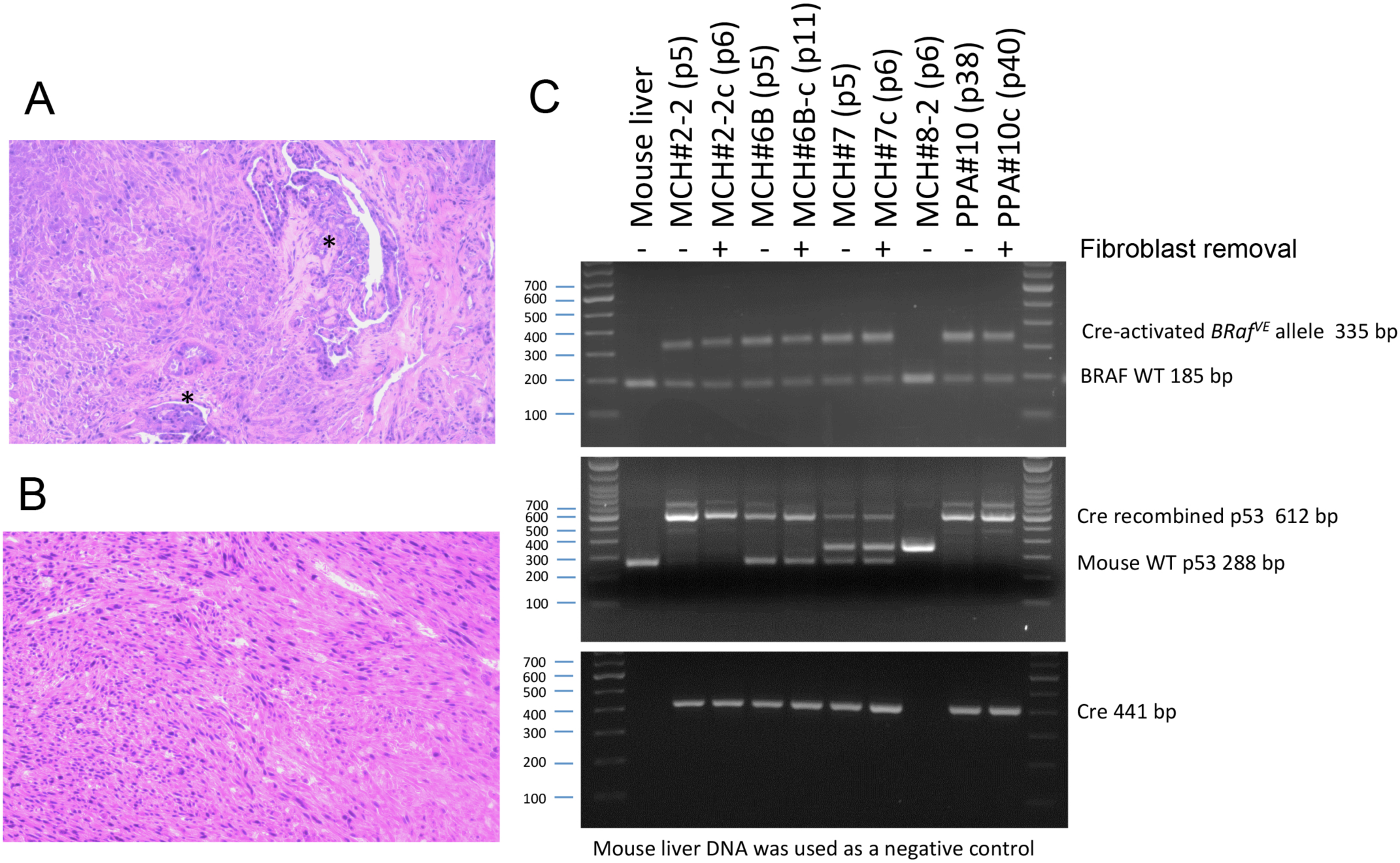
Histology of mouse ATC tumors and genotyping of derived cell lines. **A+B:** Thyroid tumors obtained from *Tpo-Cre^+^; Braf^Ca/+^; Trp53^del/del^* mice. Tumor A (PPA6) is mixed PTC/ATC with residual PTC foci indicated by asterisks (*). Tumor B (PPA10) has an entirely ATC phenotype. **C:** Genotyping of some cell lines derived from mouse thyroid tumors. MCH8-2 cells do not express *Tpo-Cre* and express only the wild-type (WT) *Braf* and *Trp53*. MCH2.2, MCH6B, MCH7 and PPA10 are all ATC cell lines heterozygous for the *BRAF^V600E^* transgene (*Ca/+*). MCH2.2 and PPA10 are homozygous for the *Trp53* mutation (exons 2-10 deletion), and XY (not shown). MCH6 and MCH7 are heterozygous for the *Trp53* mutation, and XX (not shown).

**Supplementary Figure 2:**
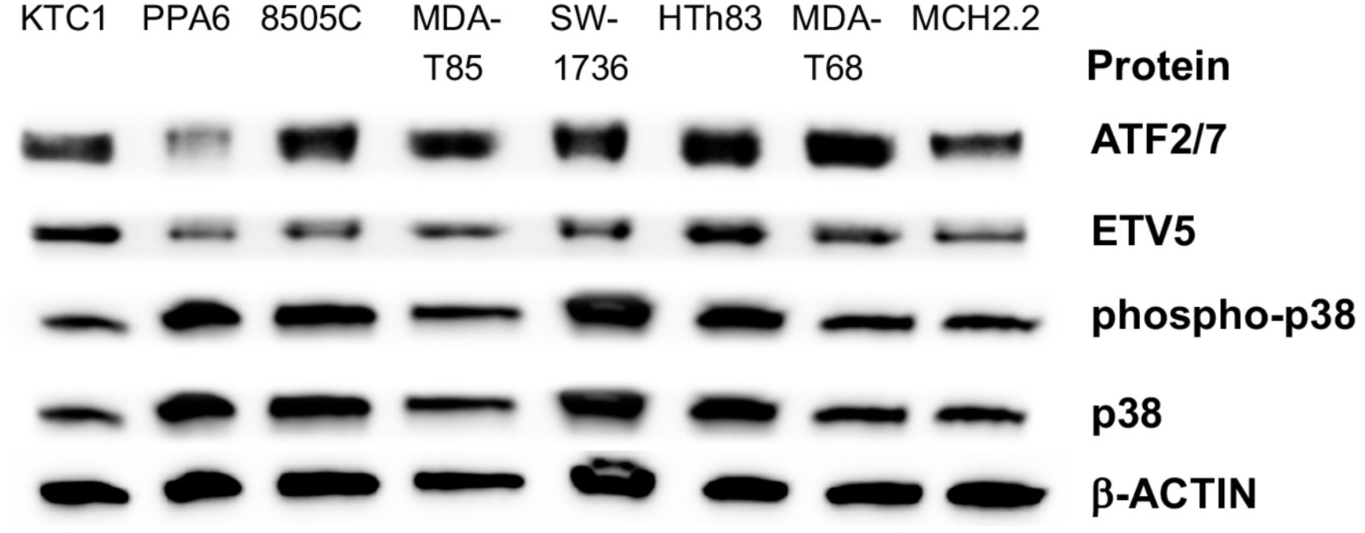
Basal protein expression of ETV5, p38, phopho-p38 and ATF2/7, a known target of activated p38, in several PDTC and ATC cell lines.

**Supplementary Figure 3.**
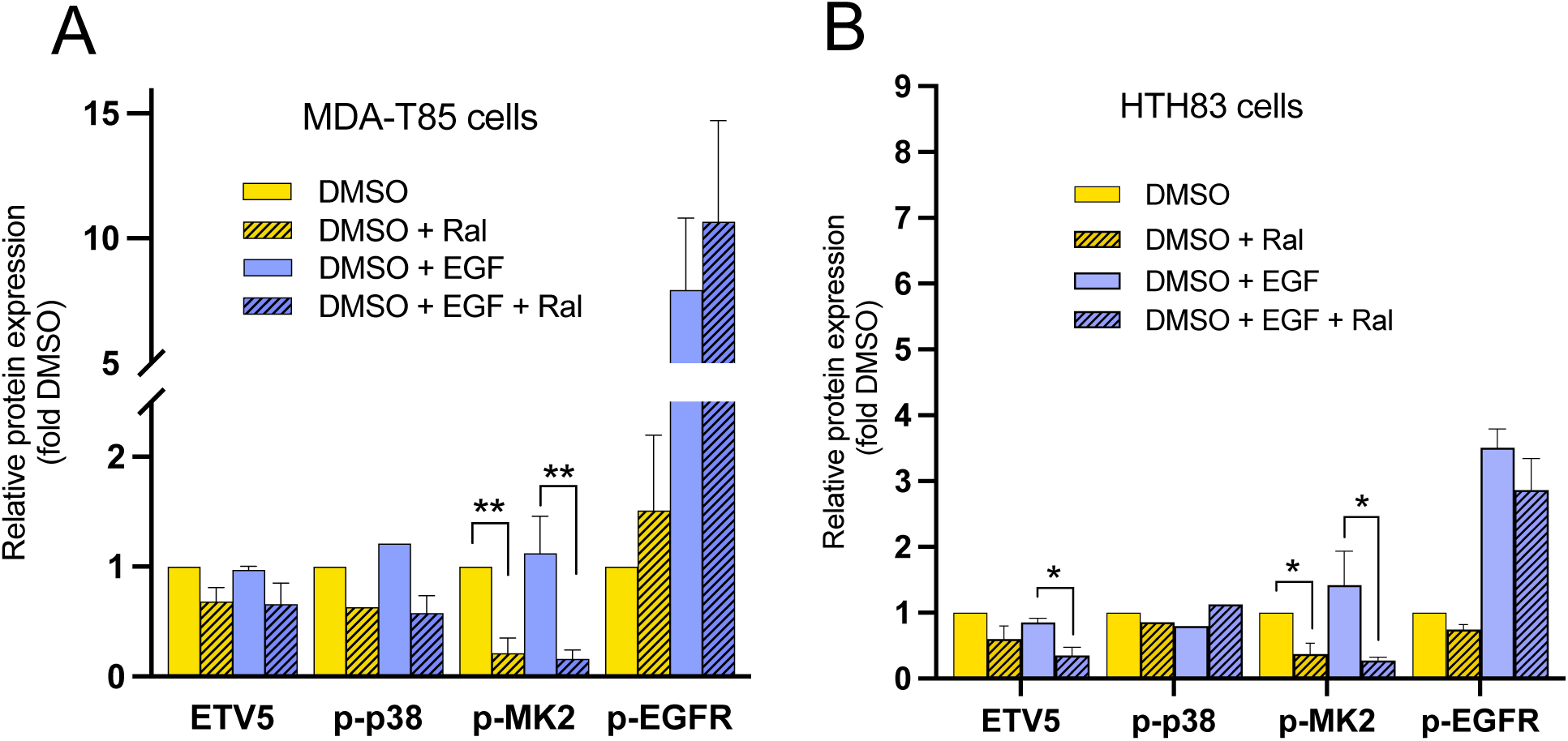
Effects of ralimetinib on FGFR and p38 pathway activity. Western blot quantification of ralimetinib effects on the phosphorylation of p38, MK2 and FGFR. **(A)** MDA-T85 (PDTC) cells and **(B)** Hth83 (ATC) cells. MK2 is a known downstream effector of p38. Cells were serum-started for 24h before adding 0.03% DMSO (ralimetinib solvent), ralimetinib, epidermal growth factor (EGF), or EGF + ralimetinib for 15 min. * = *P* < 0.05, ** = *P* < 0.01, n = 4.

## Notes

### Competing Interest Statement

Disclosures:
A Maniakas reports research funding from Thryv Therapeutics and JAZZ Pharmaceuticals
SY Lai is a medical affairs consultant for Cardinal Health
M Zafereo reports research grant funding for clinical trials from Merck, Eli Lilly, and Exelixis. ME Cabanillas reports research grants funding for clinical trials from Exelixis, Genentech, and Merck. She reports consulting Novartis, Thryv Therapeutics, Bayer, Exelixis, and Eli Lilly.
MC Hofmann reports research funding from Thryv Therapeutics
All other authors declare no potential conflicts of interest.

### Summary of Updates

Main text: readability was improved and several statements were clarified Supplementary tables: Two paragraphs were swapped and supplementary tables had to be renumbered Figures: readability of figure legends was improved/clarified and labels of some figures were edited/enlarged

