## Supplemental Table 1 for "Role of the ETV5/p38 signaling axis in aggressive thyroid cancer cells"

|  | MDA-T68  (hPTC) | MDA-T85  (hPDTC) | KTC1  (hPDTC) | KTC1-VA7  (hPDTC-ATC) | Hth83  (hATC) | SW1736  (hATC) | MCH2.2  (mATC) | PPA6  (mATC) |
| --- | --- | --- | --- | --- | --- | --- | --- | --- |
| BRAF | WT | **V600E** | **V600E** | **V600E** | WT | **V600E** | **V600E** | **V600E** |
| TERT promoter | WT | **C228T** | **C250T** | **C250T** | **C228T** | **C228T** | WT | WT |
| TP53 | WT | WT | WT | WT | **P153Afs*28** | **Q192Ter** | **Del** | **Del** |
| KRAS | WT | WT | WT | **G12D** | WT | WT | WT | WT |
| NRAS | **Q61K** | WT | WT | WT | WT | WT | WT | WT |
| HRAS | WT | **Q61K** | WT | WT | **Q61R** | WT | WT | WT |
| CDKN2A | WT | WT | **Del** | **Del** | WT | WT | WT | WT |
| AR | WT | WT | WT | WT | **Gly456_**  **Gly457insGly** | WT | WT | WT |
| TSHR | WT | WT | WT | WT | WT | **I486F** | WT | WT |

**Supplementary Table S1: Genomic landscape of the cell lines used**
