## Supplemental Table 2 for "Role of the ETV5/p38 signaling axis in aggressive thyroid cancer cells"

**Supplementary Table S2: TaqMan assays used for RT-qPCR**

| **Gene** | **Assay ID** |
| --- | --- |
| ETV5 | Hs00927557_m1 |
| TWIST1 | Hs04989912_s1 |
| SNAI1 | Hs00195591_m1 |
| GAPDH | Hs02758991_gl |
| ACTB | Hs01060665_g1 |
