## Supplemental Table 3 for "Role of the ETV5/p38 signaling axis in aggressive thyroid cancer cells"

**Supplementary Table S3**. **Primary antibodies**

| **Primary Antibody Target** | **Host/**  **Isotype** | **Dilution/**  **Concentration** | **Source** | **Catalog Number** |
| --- | --- | --- | --- | --- |
| ETV5  ETV5  p38 | Rabbit polyclonal IgG  Rabbit polyclonal IgG  Rabbit polyclonal IgG | 1:1000  1:1000  1:1000 | Abcam  ProteinTech  Cell Signaling  Technology | ab102010  13011-1-AP  8690S |
| phospho-p38  (Thr180/Tyr182) | Mouse monoclonal IgG | 1:1000 | Cell Signaling Technology | 9216S |
| Phospho-MSK1 (Thr581) | Rabbit polyclonal IgG | 1:1000 | Cell Signaling Technology | 9595 |
| Phospho-MKK3 (Ser189)/MKK6  (Ser207) | Rabbit monoclonal IgG | 1:1000 | Cell Signaling Technology | 12280 |
| Phospho-HSP27  (Ser82) | Rabbit monoclonal IgG | 1:1000 | Cell Signaling Technology | 9709 |
| Phospho-MAPKAPK-2  (Thr334) | Rabbit monoclonal IgG | 1:1000 | Cell Signaling Technology | 3007 |
| Phospho-ATF-2 (Thr71)/ATF-7  (Thr53) | Rabbit monoclonal IgG | 1:1000 | Cell Signaling Technology | 15411 |
| ERK1/2 | Rabbit polyclonal IgG | 1:1000 | Cell Signaling Technology | 4695 |
| Phospho-ERK1/2 (Thr202/Tyr204) | Rabbit polyclonal IgG | 1:1000 | Cell Signaling Technology | 4370 |
| KRAS^G12D^ | Rabbit polyclonal IgG | 1:1000 | Cell Signaling Technology | 14429 |
| STAT3 | Rabbit polyclonal IgG | 1:2000 | Cell Signaling Technology | 30835 |
| Phospho-STAT3 (Tyr705) | Rabbit polyclonal IgG | 1:1000 | Cell Signaling Technology | 9145 |
| Phospho-STAT3  (Ser727) | Rabbit polyclonal IgG | 1:1000 | Cell Signaling Technology | 9134 |
| ACTB (Actin)  Alpha-Tubulin  (TUBA1A) | Rabbit monoclonal IgG  Rabbit monoclonal IgG | 1:1000  1:1000 | Cell Signaling Technology  Cell Signaling Technology | 4970S  2144 |
| Vinculin (VCL) | Rabbit monoclonal IgG | 1:500 | Cell Signaling Technology | 13901P |
| Lamin A  GAPDH | Mouse monoclonal IgG  Rabbit monoclonal IgG | 1:1000  1:1000 | Cell Signaling Technology  Cell Signaling Technology | 86846  5174S |

**Table 4. Secondary antibodies**

| **Secondary Antibody** | **Host/**  **Isotype** | **Dilution/**  **Concentration** | **Source** | **Catalog Number** |
| --- | --- | --- | --- | --- |
| Rabbit IgG-HRP | Donkey polyclonal | 1:3000 | GE Healthcare | NA934 |
| Rabbit IgG-HRP | Donkey polyclonal | 1:3000 | GE Healthcare | NA934V |
| Rabbit IgG- HRP | Goat polyclonal | 1:5000 | Cell Signaling Technology | 7074P2 |
| Mouse IgG-HRP | Goat polyclonal | 1:2000 | GE Healthcare | NA931V |
