## Supplemental Table 4 for "Role of the ETV5/p38 signaling axis in aggressive thyroid cancer cells"

**Supplementary Table S4. IC50 values for different cell lines and pharmacological inhibitors**

*** =	positive control agents**

**# =	negative control cell lines (no BRAF mutation)**

|  | **Anisomycin*** | **Doxorubicin*** | **Dabrafenib** | **Ralimetinib** | **SB203580** | **PD169316** | **Harmine** | **Olaparib** | **Venetoclax** | **Xanthohumol** |
| --- | --- | --- | --- | --- | --- | --- | --- | --- | --- | --- |
| **MDA-T85** | **50 nM** | **10.6 nM** | **0.3 µM** | **13.3 µM** | **31.6 µM** | **31.6 µM** | **10 µM** | **3.1 µM** | **10 µM** | **12.6 µM** |
| **MDA-T68 ^#^** | **63 nM** | **10.0 nM** | **>50 µM** | **15.8 µM** | **>50 µM** | **>50µM** | **10 µM** | **3.1 µM** | **12.5 µM** | **12.5 µM** |
| **SW1736** | **ND** | **6.9 nM** | **1.3 µM** | **23.7 µM** | **>50 µM** | **>50 µM** | **31.6 µM** | **17.7 µM** | **13.4 µM** | **23.9 µM** |
| **Hth83 ^#^** | **ND** | **5.6 nM** | **31.6 µM** | **31.6 µM** | **>50 µM** | **44.6 µM** | **25.1 µM** | **13.4 µM** | **10 µM** | **7.5 µM** |
| **MCH2.2** | **31.6 nM** | **30.5 nM** | **1.0 µM** | **10.0 µM** | **10.0 µM** | **10.0 µM** | **17.7 µM** | **6.3 µM** | **3.16 µM** | **3.1 µM** |
| **PPA-6** | **0.1 µM** | **31.6 nM** | **0.2 µM** | **10.0 µM** | **15.8 µM** | **31.6 µM** | **15.8 µM** | **6.3 µM** | **3.16 µM** | **4.0 µM** |
